# The washing machine as a reservoir for transmission of ESBL-producing *Klebsiella oxytoca* in newborns

**DOI:** 10.1101/354613

**Authors:** R.M. Schmithausen, M. Exner, C. Rösing, M. Savin, S. Hack, S.F. Bloomfield, M. Kaase, J. Gebel, S. Engelhart, D. Exner

**Author notes:** Current address: Department for Infection Control and Infectious Diseases, University Medical Center Göttingen, Kreuzbergring 57, 37075 Göttingen, Germany. Corresponding author: Ricarda Schmithausen, Institute for Hygiene and Public Health, Sigmund-Freud-Str. 25, 53105 Bonn.

## Abstract

During the period from April 2012 to May 2013, 13 newborns and one child in a paediatric hospital ward in Germany were found to be colonised with a distinct clinical clone of an (extended spectrum β-lactamase) (ESBL)-producing *Klebsiella oxytoca*. This clone was specific to this hospital and had not been previously isolated in Germany.

A source-tracking analysis was carried out to identify the source and transmission pathways of the ESBL-producing *K. oxytoca* clone. A systematic environmental survey of the ward and an audit of the procedures for cleaning and disinfecting surfaces, instruments, incubators, and washing machines was performed. Microbiological samples were obtained from environmental surfaces. Risk factors were analysed for epidemiological linkage.

Isolates of an ESBL-producing *K. oxytoca* were found in the detergent drawer and on the rubber door seal of a washing machine and in two sinks. These strains were typed by pulsed-field-gel-electrophoresis (PFGE) and compared with the isolates from the newborns and their clothing and found to be identical. The retrospective analysis demonstrated that only newborns who had worn clothes that had been washed in the washing machine were colonised with the identical clone. After the washing machine was taken out of use, no further cases were detected over the following 4-year period.

We conclude that washing machines are potential reservoirs and vectors for transmission of *Enterobacteriaceae*, and likely other bacteria.

**Importance:** Washing machines should be further investigated as possible sites for horizontal gene transfer (ESBL-/carbapenemase-genes) and cross-contamination of clinically important Gram-negative strains. Particularly in the healthcare sector, the knowledge of possible (re-) contamination of laundry (patients clothes, staff uniforms) with multidrug-resistant Gram-negative bacteria could help to prevent and control nosocomial infections. To date, the potential of the washing machine as a source and vector of antibiotic-resistant gram-negative bacteria causing an outbreak in a clinical setting has not been investigated. This report shows that sampling of washing machines should be included in environmental audits associated with outbreak control management, and conditions for the laundering of baby 64 clothing should be reviewed.

## Introduction

### Background

Newborns are vulnerable to sporadic infections and outbreaks caused by nosocomial pathogens. Frequent reports have described potential risk factors associated with multidrug-resistant *Klebsiella* spp. (1–9). Colonisation with antibiotic-resistant bacteria such as ESBL-producing *Klebsiella oxytoca* can lead to clinical disease in newborns, with life-threatening consequences (10–15). Nevertheless, the causal environmental reservoir and transmission pathways have been elucidated in only a few outbreak studies (16–18).

For the sustainable control of outbreaks, it is imperative to elucidate the causal reservoir and bring it under control. The new 2017 German legislation demands that probable infection reservoirs must be identified by typing methods as a result of source tracking. Washing machines and clothing are not currently assessed as potential reservoirs and transmitters in outbreaks of nosocomial infections, despite evidence that a potential health risk due to cross-contamination of contaminated laundry cannot be excluded (19–21).

### Case description

During the period from April 2012 to May 2013, increased rates of colonisation (no infection) with ESBL-producing *Enterobacteriaceae*, particularly *K. oxytoca* isolates, were recorded in a perinatal centre (PNC) and on several wards in a children’s hospital in Germany. Isolates were considered as hospital-acquired if the first specimen yielding resistant *K. oxytoca* was obtained ≥ 3 days following admission or if the specimen was obtained < 3 days following admission in a patient who had been hospitalised at the outbreak hospital within the previous 3 months. Patients were characterised as colonised but not infected according to CDC definitions (22). The first seven cases were identified following the introduction of a standard screening procedure. During the period from August to October 2012, *K. oxytoca* was repeatedly identified in 5 out of 80 infants born in the Obstetrics Ward and treated in the PNC.

## Methods

### Control management and containment measures

When the first cases occurred in April 2012, the responsible clinician informed the Head of the Infection Control Department and the Public Health Officer of the Public Health Department, and a task force was constructed. The task force developed a targeted management strategy to analyse the cases, while also focusing on common risk factors to identify the source/reservoir of the *K. oxytoca* strain and allow the outbreak to be controlled. The detailed management procedures are presented in Table S1.

Within three management phases, a screening of all healthcare workers at the PNC, Intensive Care Unit (ICU), and Obstetrics Department was obtained. However, since no sources or transmission route had been identified and positive cases were still occurring following two rounds of infection control measurements, by October 2012, a third phase of measures (October 2012 to June 2013) was initiated (Table S1). In March 2013, cases were still occurring and it was decided to consult an infection control expert who reviewed the measures taken and recommended identification and sampling of specific risk factors (water/wastewater reservoirs, e.g. washing machines).

### Screening of the newborns, mothers, and healthcare workers

All paediatric patients from six different wards (including the ICU and the PNC) were screened for ESBL-producing *Enterobacteriaceae*. Screening swabs were taken from the anal region, throat/sputum, and wounds. Newborns were screened directly after birth, 48 hours after birth, and weekly thereafter. All other inpatient children were screened at admission and weekly thereafter. In addition, an intensified screening of all mothers, firstly on admission and secondly on discharge, was performed to rule out nosocomial contamination during their stay.

Further screening of all healthcare workers (physicians, nurses, and cleaning personal working in the PNC, ICU, and Obstetrics Department) was conducted to identify any carriers of the *K. oxytoca* strain.

### On-site inspections and environmental monitoring

Several on-site inspections were made in all areas that were functionally related to the affected wards. The first inspection was performed in October 2012. Samples of surrounding areas were obtained in May and October 2012 by the intern Infection Control Expert and by the Head of the Microbiology Laboratory. The second on-site inspection focusing on the PNC took place in June 2013 by an external Hygienist together with the Public Health Officer. Environmental and water samples were taken by an experienced Infection Control Practitioner. An overview of the analyses of the first and second on-site inspections and related measures is given in Table S2. A detailed list of the sampling points and types is given in the Supplemental Material and Table S3.

During the first on-site inspection, sampling via contact culture plates was performed in the immediate surroundings of the newborns, children, and newly admitted mothers in the PNC, ICU, and Obstetrics Department. During the second on-site inspection, the sample area was enlarged. All premises (staff room, patient rooms, preparation rooms, and storage rooms) were sampled, including storage rooms for the preparation of the incubators and the laundry room on the ground floor.

Additional environmental samples were collected from syphons, sinks, toilets, showers, and water boilers (drinking water, wastewater, etc.) (Tables S2 and S3). All disinfectants (cloth dispenser system, etc.) and wipes for disinfecting the incubators as well as care emulsions from staff and patient rooms were collected for laboratory analyses (Table S3). Swab samples were obtained from the two washing machines and the tumble dryer via sterile environmental swab tubes. Wastewater and residual water from the outlet drains was collected, and swabs were taken from the rubber mantle of the washing machine as well as native water samples (Table S3).

In a further step, all clothing that had been washed in one of the washing machines and worn by the affected newborns was collected and microbiologically analysed. These were mainly socks and hats knitted from cotton and wool. No documentation regarding the distribution of responsibilities or manner of use of the washing machine exists; therefore, the temperature at which clothing worn by newborns and other laundry were washed in this washing machine is untraceable. According to protocol, all clothing should have been washed at a temperature of at least 65°C in a “medical wash” program; however, as far as can be reconstructed, in this case, the socks and hats had been washed at 40°C with an approved industrial disinfectant detergent.

### Laboratory analyses

Environmental samples obtained during the first on-site inspection were analysed in a laboratory at the Bionovis Hygiene Institute Joachim Kruff e.K. in Giessen. Environmental probes from the second on-site inspection were analysed at the Institute for Hygiene and Public Health of the University Hospital, Bonn. All staff and newborn samples were analysed in the Microbiology Laboratory of the affected hospital (Laboratory Corporation, Dr. Dirkes-Kersting and Dr. Kirchner mbH, Siegen). All samples were stored at 4°C during transport to the laboratories and inoculated within 48 hours.

All human samples were streaked on MacConkey (Oxoid Deutschland GmbH, Wesel, Germany) with a 10-μg imipenem disk (bioMérieux SA, Marcy-Étoile, France) and selective agar plates, i.e. CHROMagarESBL (Oxoid Deutschland GmbH, Wesel, Germany) with a 10-μg ertapenem disk (bioMérieux SA).

At the Bionovis Hygiene Institute, all BD RODAC^TM^ contact plates (∅ 55 mm, Becton & Dickinson, Heidelberg, Germany) were incubated at 36 ± 1°C for 48 hours. The swab specimen was spread onto MacConkey (Oxoid Deutschland GmbH, Wesel, Germany) and BD Columbia Agar with 5% sheep blood.

At the Institute of Hygiene and Public Health in Bonn, all environmental swabs were streaked on Columbia / 5% sheep blood agar plates (Becton Dickinson, Heidelberg, Germany), MacConkey (Oxoid Deutschland GmbH, Wesel, Germany; Art. No. PO5002A), and selective agar plates, i.e. CHROMagarESBL (bioMérieux SA, Marcy-Étoile, France; Art. No. 43481). Plates were incubated at 37 ± 1°C for 48 hours. Additionally, the swabs were inoculated into Caseinpepton-Soy flour peptone-bouillon (CASO-bouillon) (Merck KGaA, Darmstadt, Germany). The CASO-bouillons were incubated at 37 ± 1°C for 24 hours.

The disinfectant wipes, socks, and knitted hats of the newborns were inoculated into 200 mL CASO-bouillon with TSHC (+3% Tween Merck, Art. No.: 8.22187; +3% Saponine, Roth, Art. No. 5185.1; +0.1% Histidine Merck, Art. No. 1.04351; +0.1% Cysteine, Merck, Art. No. 1.02839; TSHC) (Art. No. 1.05459, Merck KGaA, Darmstadt, Germany), homogenised for 60 seconds (Stomacher 400, Seward Limited, West Sussex, United Kingdom), and filtered (membrane-filter Millipore Hydrosol, 250 ml cup). 2 × 100 mL extract was filtered on ESBL and MacConkey agar, rinsed with 100 mL 0.9% NaCl, and incubated at 37°C for 24–48 hours. In parallel, a dilution series was made, and 100 μL of each concentration was plated on both ESBL and MacConkey agar. The emulsions were treated in the same manner as the wipes, except that shredding was not necessary.

For the water samples, approximately 100 mL was collected in a sterile polystyrene cup (article no. 225170, Greiner, Frickenhausen, Germany) and filtered twice (2 × 50 mL) through a sterile nitrocellulose membrane filter (pore size 0.45 ± 0.02 μm, ∅ 47 mm, black grid from Millipore, Art. No. EZHAWG 474), according to Schulz and Hartung (2009). After filtration, the membrane was placed on selective CHROMagar ESBL and MC agar plates (bioMérieux SA, Marcy-Étoile, France). In addition, three 10-fold dilutions were prepared using sterile NaCl, and 100 μL each dilution was plated on CHROMagar ESBL and MC agar.

All *Enterobacteriaceae* detected on CHROMagarESBL were identified via API 20 E V4.1 apiweb^™^ (bioMérieux SA, Marcy-Étoile, France) and matrix-assisted laser desorption/ionisation time-of-flight mass spectrometry (MALDI-TOF MS, bioMérieux SA or Bruker, Billerica, USA).

Antibiotic susceptibility was determined on a VITEK-2 (bioMérieux SA), and results were interpreted using EUCAST criteria. Susceptibility to antibiotics was tested using the Micronaut-S MDR MRGN-Screening 3 system (MERLIN, Gesellschaft für mikrobiologische Diagnostika GmbH, Bornheim-Hersel, Germany).

ESBL-production was confirmed by PCR detection of CTX-M. To this end, three specific primer sets were used to detect β-lactamase-encoding genes belonging to the *bla*TEM, *bla*SHV, and *bla*CTX-M families (23–25). For DNA template preparation, one loop of bacteria (1 μL) was resuspended in 200 μL 10 mM Tris-EDTA buffer (pH 8) and incubated for 10 minutes at 95°C. Cell debris was removed by centrifugation (20,000 × g for 3 minutes). The following PCR conditions were employed: a 4-minute initial denaturation at 94°C; 35 cycles of denaturation at 94°C (30 seconds), annealing at 55°C (30 seconds), elongation at 72°C (60 seconds); followed by a final extension at 72°C for 5 minutes. The PCR products were visualised by gel electrophoresis on a 1% agarose/TBE gel, and stained with Midori Green (Biozym Scientific GmbH, Hessisch Oldendorf, Germany). The resulting amplicons were purified using an innuPREP DOUBLEpure Kit (Analytik Jena AG, Jena Germany) according to the manufacturer's recommendations. Custom-sequencing was performed by Microsynth (Göttingen, Germany). The nucleotide sequences were analysed using Chromas 2.6.5.

All *K. oxytoca* isolated from clinical samples and environmental sources were typed by the German National Reference Centre (NRC) using pulsed-field gel-electrophoresis (PFGE), according to Tenover et al., (1995) (26) and modified according to Ribot et al., (2006) (27).

## Results

### Detailed outbreak course and results of the clinical and staff samples

During the period from April 2012 to May 2013, a total of 27 children were colonised with *K. oxytoca*. Almost every month, one additional case was identified. The occurrence of *K. oxytoca* varied from sensitive isolates with no ESBL-activity to ESBL-producing *K. oxytoca* and the particular ESBL-producing *K. oxytoca* type-00531. Subtyping shows that the *K. oxytoca* type-00531 has never been detected before in the NRC. According to the numbering of the NRC, the term “special *K. oxytoca* “type-00531” was used for description.

The ESBL-producing *K. oxytoca* strains were intermediately resistant to piperacillin/tazobactam, with a minimum inhibitory concentration (MIC) of 16 mg/L, and resistant to cefotaxime (MIC ≥ 64 mg/L) and ceftazidime (MIC 8 mg/L). Considering this, according to the German Classification for Multidrug-resistant Gram-negative Microorganisms (MRGN), the strains could be classified as “2MRGN NeoPäd”.

Fig. 1 summarises the occurrence of cluster-related ESBL-producing *K. oxytoca* clinical isolates over time in the PNC, ICU, and on four different wards. Fourteen children were positive for the ESBL-*K. oxytoca* type-00531. Four *K. oxytoca* strains were carrying ESBL but could not be classified as type-00531. In nine children, *K. oxytoca* without the ESBL-enzyme was detected.

**Figure 1.**
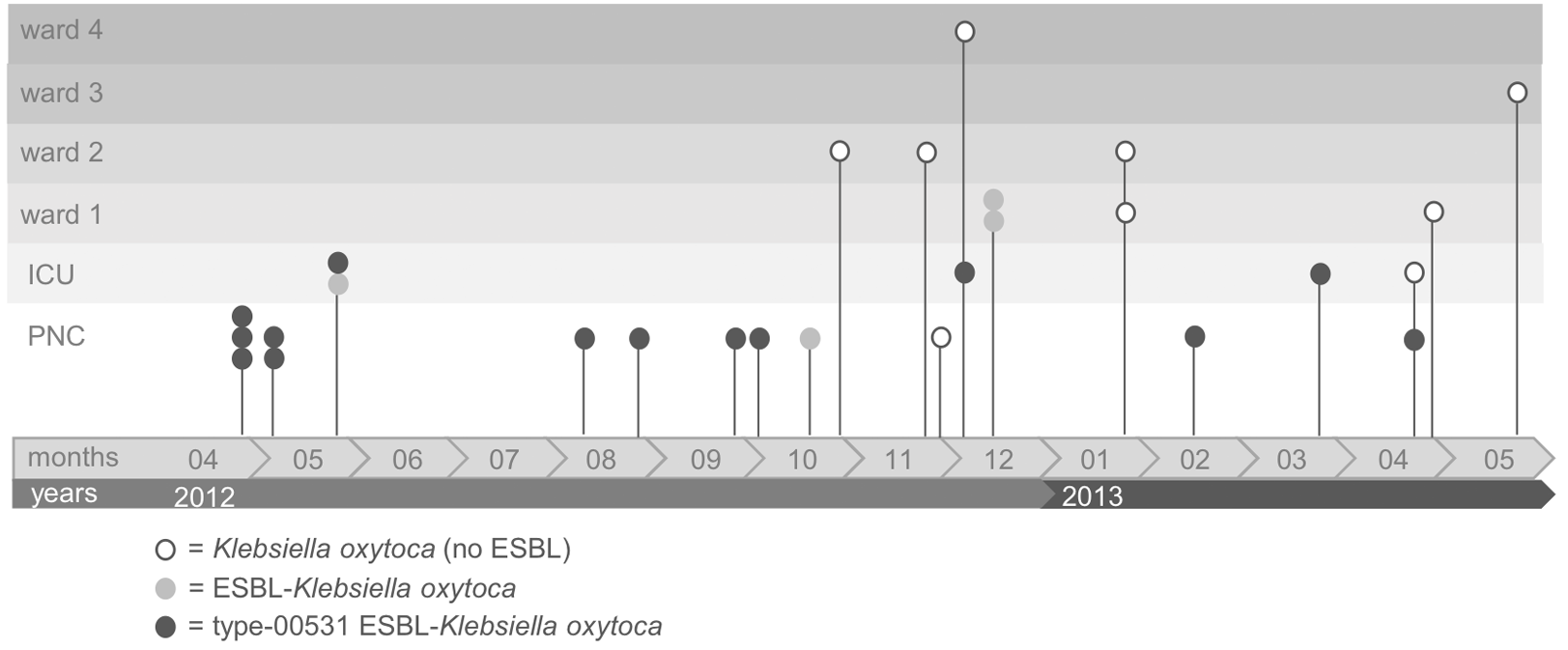
Course of outbreak with different *K. oxytoca* strains within one year and their distribution on different wards (PNC: perinatal centre, ICU: intensive care unit).

Fig. 2 shows the age distribution of the colonised children over the entire outbreak period. For all newborns and infants, *K. oxytoca* was mostly identified by anal swab screening. In 9 out of 24 newborns (1-4 weeks old) *K. oxytoca* was also detected in throat samples. The older children were more likely to have colonised wounds (no ESBL-*K. oxytoca*, 14 years old, ward 4) or PEG-tubes (percutaneous endoscopic gastrostomy) (no ESBL-Klebsiella, 17 years old, ward 3). However, a 4-year-old Arabic boy in the ICU, who never had any contact with the PNC, also showed anal colonisation with type-00531 ESBL-*K. oxytoca* in March 2013.

**Figure 2.**
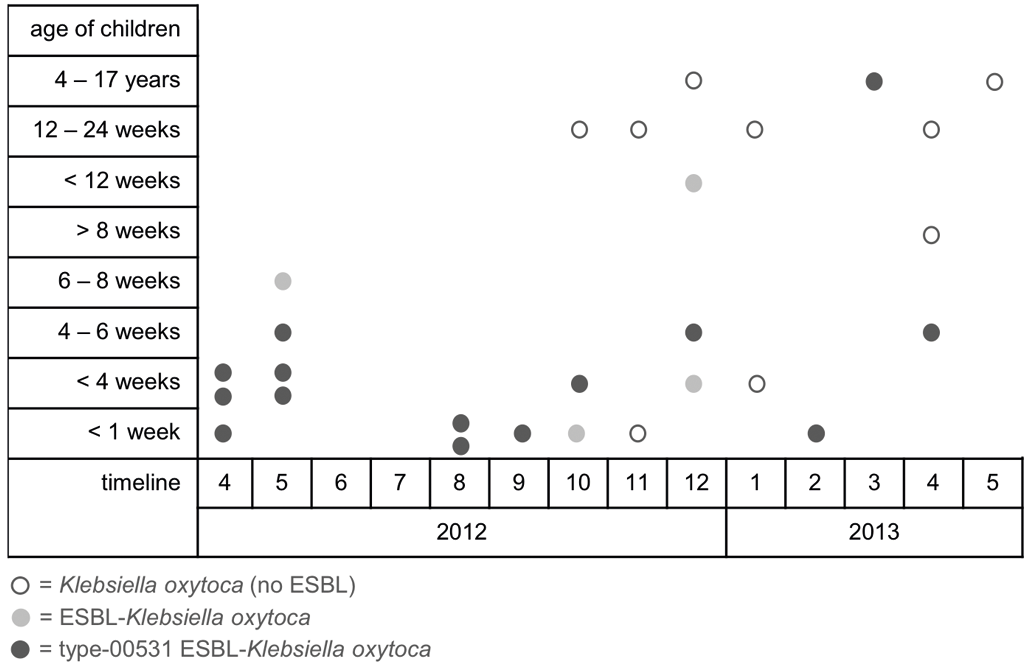
Distribution of different *K. oxytoca* strains over the “outbreak period” regarding the age of newborns and children.

Until October 2012, only newborns (< 4 weeks old) in the PNC or infants (6-8 weeks old) in the ICU were affected with ESBL-*K. oxytoca*, either with or without type-00531 (Fig. 1. From the end of October 2012 to February 2013, *K. oxytoca* was also detected on other wards and in older paediatric patients (see above); however, these strains were not similar to the type-00531 that had been prominent in the previous months.

Among these was a newborn (aged <4 weeks) who initially tested negative in the PNC, but subsequently tested positive for ESBL-*K. oxytoca* (nontype-00531) during the screening procedure in the ICU. From August 2012 to March 2013, a total of six cases were positive for ESBL-*K. oxytoca* type-00531, with 5 of the 6 cases (4 caesarean sections, 2 vaginal deliveries) being positive in the second testing but negative in the initial screening (postpartum).

During an extended screening of mothers and healthcare workers, a total of 695 swabs from 428 persons (vaginal and rectal) were obtained on the Obstetrics Ward. The screening identified four cases of *K. oxytoca* with typical resistance, five cases with ESBL-producing *E. coli*, and 1 ESBL-producing *K. pneumoniae.* Although most of the mothers were treated with antibiotics, no case of transmission between mother and newborn was documented. No person was positive for the special type-00531 ESBL-producing *K. oxytoca*.

### Results of the environmental monitoring

All environmental samples (Tables S2 and S3) taken during the first and second on-site inspections tested negative for Gram-negative *Enterobacteriaceae* and non-fermenting organisms. During the second on-site sampling in October 2012, only low concentrations of Gram-positive skin and environmental bacteria were detected: *coagulase-negative staphylococci*, *micrococci*, and gram-positive rods. However, during the third on-site inspection in June 2013, Gram-negative *Enterobacteriaceae* and non-fermenting organisms were detected. All *K. oxytoca* isolates were identical ESBL-producing strains and belonged to type-00531 (Table 1).

**Table 1.**
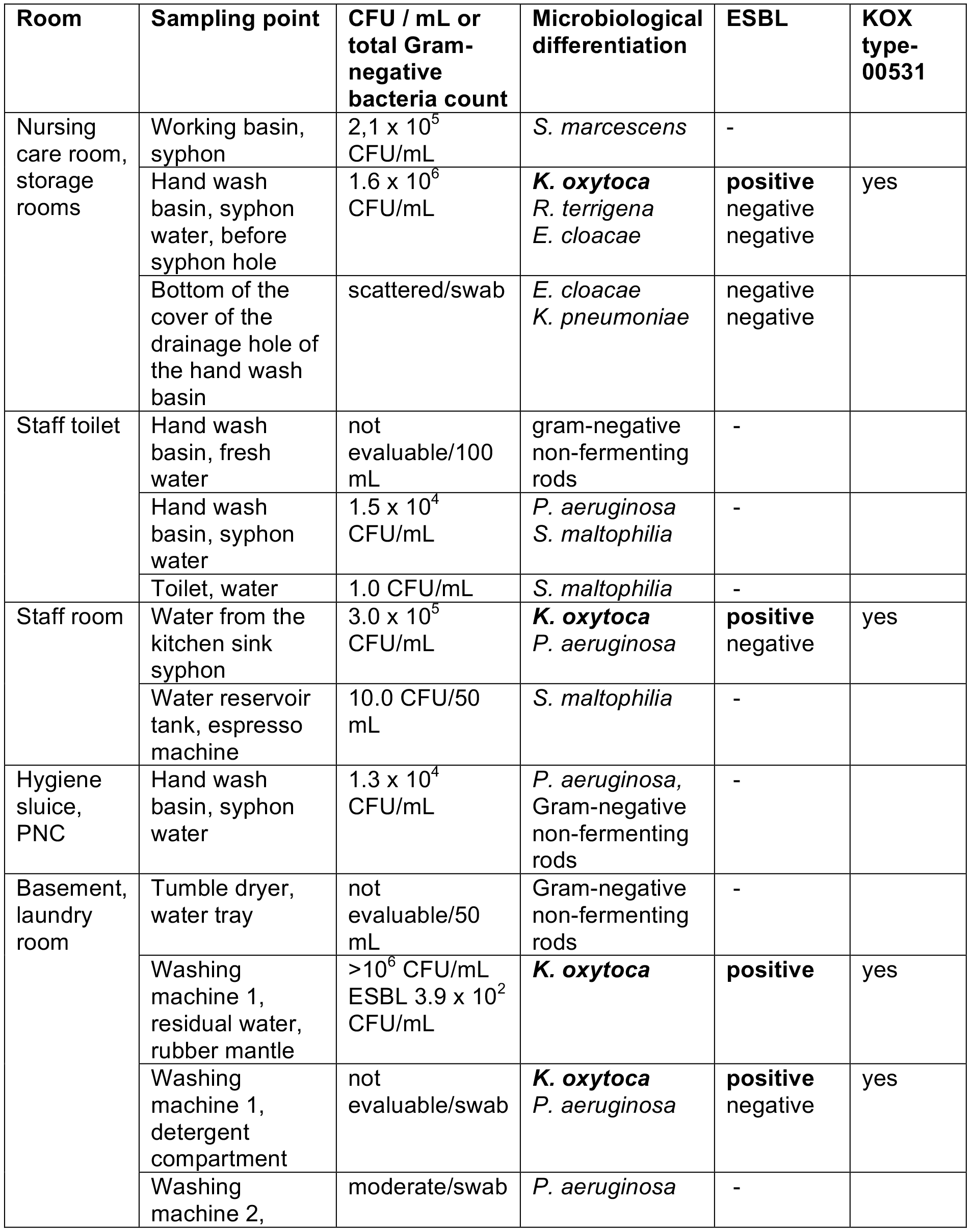
Occurrence of Gram-negative *Enterobacteriaceae*, non-fermenting organisms, and 629 ESBL-producing *Klebsiella oxytoca* type-00531 isolates in environmental samples obtained 630 during on-site inspection of risk areas.

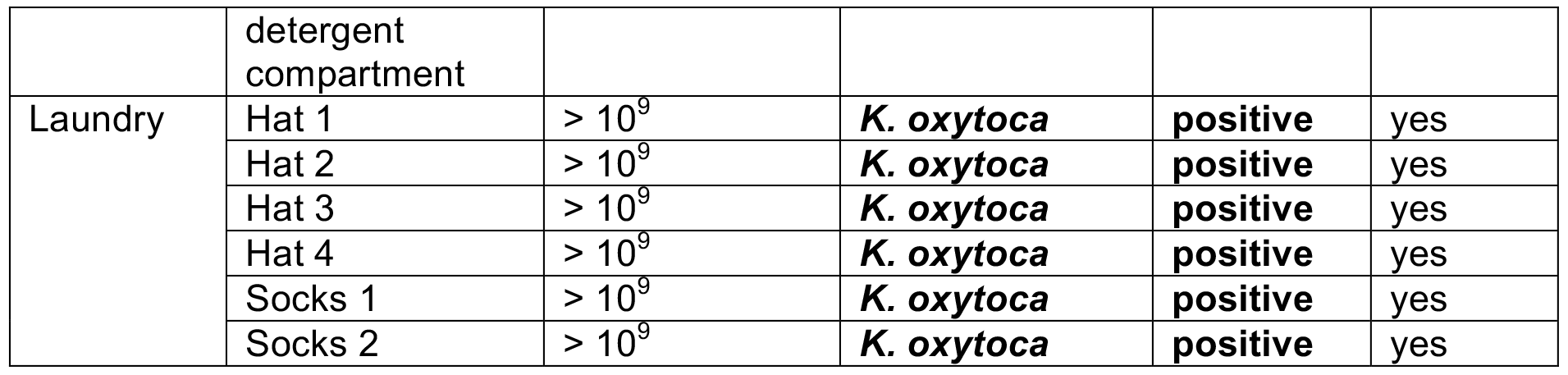

Water-associated bacteria such as *P. aeruginosa*, *Serratia* spp., *Enterobacter* spp., *K. pneumoniae*, and *Stenotrophomonas maltophilia* were detected in the syphons of hand wash basins. Identical clones of *K. oxytoca* type-00531 were isolated from the syphons of two sinks in the healthcare workers’ staff room and in the room used for cleaning and disinfection. *K. oxytoca* could be detected prior to but not following water flushing of the sinks. Accordingly, following periods of stagnation, a significant increase in the level of contamination in the syphon water up to 10^6^ CFU / mL was observed. The same clone was also isolated in high concentrations (Table 1) from samples of residual water in the rubber seal and one swab sample (in addition to *P. aeruginosa*) from the detergent compartment of one of the two washing machines.

Following identification of the washing machine as a potential reservoir for this specific *K. oxytoca* clone, newborn clothing (hats and socks) that had been washed in this machine were microbiologically analysed. *K. oxytoca* of the same specific clone was isolated with a total count of > 10^9^.

All ESBL-producing *K. oxytoca* type-00531 found in the water and environmental samples were also intermediately resistant to piperacillin/tazobactam, with a minimum inhibitory concentration (MIC) of 16 mg/L, and resistant to cefotaxime (MIC ≥ 64 mg/L) and ceftazidime (MIC 8 mg/L). Thus, since these strains were coherent to the resistance status of the newborn strains, the environmental strains could also be classified as “2MRGN NeoPäd”. The presence of ESBL genes was confirmed by PCR in two strains, both of which were CTX-M-15-positive.

### Epidemiological links

All clinical and environmental isolates displayed an identical banding pattern by pulsed-field gel-electrophoresis, and were thus considered clonally identical; the unique ESBL-producing *K. oxytoca* type-00531. Therefore, this distinct clinical clone was specific for the newborns/infants and the environment in this children’s hospital.

Retrospective analysis demonstrates that only newborns who had worn clothing that had been washed in the washing machine were colonised with the identical *K. oxytoca* clone. Although the syphons of the staff sinks were also identified as a possible reservoir, no staff members were identified as positive carriers or spreaders of ESBL-*Enterobacteriaceae* at the time of screening.

### Long-term clinical effects / concomitant infection control interventions

After the washing machine was taken out of use, no further cases of colonisation of newborns by *K. oxytoca* have been detected to date. All washing machines used for internal cleaning of patient-associated clothing were removed, and thereafter, all garments worn by the newborns and children were laundered by a professional external hospital laundry service. The two colonised sinks were replaced by new sinks with specialised thermosyphon systems. As further consequences of the prolonged cluster process, the existing infection control measures (isolating colonised patients, enforcing hand hygiene measures, and cleaning the ward, particularly the sinks and equipment) were reinforced. The screening plan for all newborns and children, and all measures implemented within the containment program, have been maintained to date.

## Discussion

### Waterborne outbreaks in healthcare systems

In hospitals, water or wastewater that is contaminated with healthcare-associated pathogens provides a potential reservoir for infections (28). Decker et al., (2013) (29) concluded that the nature of the hospital environment fosters contamination with waterborne pathogens. The most common waterborne pathogens causing healthcare-associated infections linked to contaminated hospital water are Gram-negative bacteria including *Pseudomonas* spp., *Enterobacter* spp., *Serratia* spp., *Stenotrophomonas* spp. and *Klebsiella* spp. Gram-negative bacteria are also known to pose a severe threat to high-risk patients, especially those who are immunocompromised, including severely ill patients and premature infants in the NICU (30). It is therefore unsurprising, that healthcare-associated severe infections in such vulnerable patient populations caused by *K. oxytoca* have most often been associated with contamination of environmental reservoirs (12, 16, 31–33).

In particular, Leitner et al., (2015) (34) found that multidrug-resistant organisms were most commonly linked to contaminated sinks as a reservoir, and cited an outbreak of six infections due to KPC-2-producing *K. oxytoca* in patients with haematological malignancies. However, outbreaks caused by *K. oxytoca* in neonatal care units linked to environmental sources have been rarely described (10, 11). To the best of our knowledge, the present case is the first report of a cluster of contaminations with *K. oxytoca* in newborns in a neonatal care unit linked to contaminated water or wastewater.

Although common water reservoirs in healthcare settings have been identified as tap water, faucets, sink surfaces, bathtubs, and wastewater drainage system drains, sinks, showers, toilets, and drainage pipes (30, 35), a potential reservoir that has so far been neglected is the washing machine. Only recently did Rehberg et al., (2017) (36) describe the presence of antibiotic-resistant bacteria and their possible transmission in washing machines.

### Reservoirs and transmission routes of waterborne pathogens in healthcare systems

When the first five cases occurred between August and October 2012, it was suggested that person-to-person (between patient-patient, mother-patient, or healthcare worker (HCW)-patient) transmission may have occurred, even though this has never been described for *K. oxytoca* to the best of our knowledge. Price et al., (2017) (37) showed that in the presence of standard infection control measures, HCWs were frequent sources of transmission of *S. aureus* to patients, but, thus far, transmission of Gram-negative bacteria has only been observed between patients, not originating from HCWs (38).

The screening of HCWs and mothers identified no cases of colonisation with the type-00531 ESBL-producing *K. oxytoca*. The relationship between sensitive *K. oxytoca* cases in the HCWs and the affected newborns remains unclear. An epidemiological link could neither be confirmed nor ruled out. Since the occurrence of *K. oxytoca* isolates continued over a 1-year period, a reservoir in the environment rather than hospital personnel or mothers was suspected. Despite the strict implementation of control management and containment measures, newborns and children continued to acquire the cluster organism (type-00531) and other sensitive *K. oxytoca* and ESBL-producing *Enterobacteriaceae*. The methods of sampling were changed to native water sampling and taking swabs (Table S3). It was reasoned that, although in many hygienic environmental analyses RODAC plates are applied to surfaces (39), native samples often give more profound insight. Here, the possible gain in knowledge needs to be counterbalanced against the increased efforts required in sample collection and processing in the laboratory.

### Sinks

Within the third round of environmental screening, the cluster strain was found in two sinks; in water from the kitchen sink syphon in the staff room and in water from the hand wash basin syphon in the nursing care room / storage room. Similar to the *K. oxytoca* outbreak described by Lowe et al., (2012) (16), distribution of the *K. oxytoca* strain from both sinks and colonised persons (HCWs) cannot be excluded. Sink drains are critical reservoirs, especially for Gram-negative bacteria (16, 40–43). Once colonised with *Enterobacteriaceae*, further dispersal occurs along the wastewater drainage system to distal sink drains connected to the primary reservoir. Further transmission of these bacteria from a water reservoir may occur by direct and indirect contact, ingestion, aspiration, or inhalation of aerosols (30, 35, 44). Doring et al., (1996) (40) showed that splashes of contaminated sink water could result in the colonisation of HCWs who use the sink to wash their hands, and also other items in the environment in close proximity. To prevent contamination, separate sinks should be used for handwashing and disposal of contaminated fluids (28, 44). In this case, the contaminated sinks and washbasins were removed and replaced.

### Washing machine

In this case, the particular *K. oxytoca* strain was not only detected in two sinks but also in a previously unreported water-associated reservoir; the washing machine. Even recently published articles reviewing the main water-associated reservoirs in hospitals do not consider the washing machine as a potential hazard in a clinical environment (30, 35). Thus far, case reports regarding washing machines have only focused on Gram-positive pathogens. Sasahara et al., (2010) (45) described an association between contamination of washing machines, bed linens and an outbreak of *Bacillus cereus* bacteremia. Moreover, Yoh et al., (2010) (46) discussed whether industrial laundry processes are ineffective in reducing *B. cereus* contamination of hospital linens. Furthermore, Heudorf et al., (2017) (47) mentioned that pathogens classified by the WHO as highest priority can be found on staff gowns can be the starting point of transmissions (WHO, 2017).

Recently, a few studies have suggested the potential role of washing machines in the distribution of antibiotic-resistant Gram-negative bacteria during laundering (36); however, to date, no transmission of pathogens from a washing machine to patients could be proven. In this case, we hypothesise that the *K. oxytoca* strain type-00531 was disseminated after the washing process via the residual water on the rubber mantle and/or via the final rinsing process that runs unheated and untreated water through the detergent compartment. Consequently, we conclude that newborns were colonised by wearing hats and socks that were contaminated by the washing process. The contamination of the sinks is thought to be due to the handwashing of the HCWs and not the primary source.

Finally, it remains unclear how the washing machine could become contaminated and serve as a distributor; however, once contaminated, *Enterobacteriaceae* can survive in wet environments for an extended period of time (44), especially waterborne bacteria such as *P. aeruginosa* and *Klebsiella* strains, which have the ability to survive in a viable but not culturable state. Similarly, their environmental stability is supported by the formation of biofilms to enhance their chances of multiplication and horizontal gene transfer (48).

Rehberg et al., (2017) (36) demonstrated that antibiotic-resistant bacteria can survive the washing process. For tests with *P. aeruginosa* outbreak strains, even at temperatures above 50°C, no reduction could be achieved. Even if high laundering temperatures had been used, it is likely that the temperature in the area of the rubber mantle or the rubber door seal would have been much lower, providing an optimal humid environment and nutrient supply for growth of Gram-negative microorganisms (48, 49). Moreover, the occasional use of washing machines at low temperatures supports the formation of biofilms (19).

## Conclusion

While previous studies have implicated sinks as potential reservoirs for clusters of infection caused by *K. oxytoca*, this is the first report implicating washing machines as a confirmed reservoir. Generally, to contain such a nosocomial colonisation cluster with several potential environmental reservoirs, a multi-dimensional containment approach is necessary. In this case, the approach included reinforcement of infection control policies (hand hygiene, contact precautions, isolation, admission/routine anal screening, clear delineation between handwashing sinks and sinks used for other purposes), and intensified cleaning or replacement of the sinks. However, it was only after the main environmental source, the washing machine, was removed and alternative arrangements for laundering of clothing were made, that the colonisation cluster was terminated. Based on this, washing machines should be further considered as a “playground” for horizontal gene transfer (ESBL-/carbapenemase-genes) and cross-contamination of clinically important Gram-negative strains. The results also suggest that for the prompt management of outbreaks or colonisation clusters, the choice of environmental sampling points should take into account the ecological properties of the causative strain in order to indicate where it is most likely to be found. The study also suggests that manufacturers of washing machines used in institutional settings where vulnerable patients are being cared for, should find solutions to control the growth of nosocomial pathogens. Changes to machine design and processing are required to prevent accumulation of residual water in areas of the machine, such as the detergent filling box and rubber door seals, where microbial growth can occur and contaminate clothes during the rinse cycles after they have been disinfected by laundering.

In summary, the present study shows that in situations where an increase in colonisation by species of *Enterobacteriaceae* is observed in newborns and adult patients, washing machines and clothes should be assessed and investigated as potential reservoirs and vectors for transmission.

## Acknowledgements

The authors thankfully acknowledge the consent of the concerned hospital to publish. We thank the “Laborbetriebsgesellschaft Dr. Dirkes-Kersting und Dr. Kirchner mbH, Labor Siegen” and the “Bionovis Hygieneinstitut Joachim Kruff e.K.” We gratefully acknowledge David Wellen for his active support and expert advice. We also thank Thomas Meckel for giving support regarding environmental sampling.

Special thanks to Gabi Bierbaum, Esther Sib and Gero Wilbring for their constant support during report period.

